# Long-term dynamic changes of NMDA receptors following an excitotoxic challenge

**DOI:** 10.1101/2022.01.03.474836

**Authors:** Alberto Granzotto, Marco d’Aurora, Manuela Bomba, Valentina Gatta, Marco Onofrj, Stefano L. Sensi

## Abstract

Excitotoxicity is a form of neuronal death characterized by the sustained activation of N-methyl-D-aspartate receptors (NMDARs) triggered by the excitatory neurotransmitter glutamate. NADPH-diaphorase neurons [also known as nNOS (+) neurons] are a subpopulation of aspiny interneurons, largely spared following excitotoxic challenges. Unlike nNOS (-) cells, nNOS (+) neurons fail to generate reactive oxygen species in response to NMDAR activation, a key divergent step in the excitotoxic cascade. However, additional mechanisms underlying the reduced vulnerability of nNOS (+) neurons to NMDAR-driven neuronal death have not been explored. Using functional, genetic, and molecular analysis in striatal cultures, we demonstrate that nNOS (+) neurons possess distinct NMDAR properties. These specific features are primarily driven by the peculiar redox milieu of this subpopulation. In addition, we found that nNOS (+) neurons exposed to a pharmacological maneuver set to mimic chronic excitotoxicity alter their responses to NMDAR-mediated challenges. These findings suggest the presence of mechanisms providing long-term dynamic regulation of NMDARs that can have critical implications in neurotoxic settings.

## Introduction

N-methyl-D-aspartate receptors (NMDARs) are ionotropic glutamatergic receptors primarily permeable to calcium ions (Ca^2+^). Transient NMDAR-driven Ca^2+^ influx mediates essential physiological or pathological functions (Choi, 1988, 2020; Paoletti et al., 2013). In pathological conditions, NMDAR overstimulation generates a Ca^2+^-dependent cascade of events encompassing the production of reactive oxygen and nitrogen species (ROS and RNS, respectively), irreversible mitochondrial failure, and zinc (Zn^2+^) mobilization, ultimately leading to neuronal demise (Choi, 2020; Sensi et al., 2009; Wang and Swanson, 2020). The process, termed excitotoxicity (Choi, 1992, 2020; Lau and Tymianski, 2010), is critical for the development of acute or chronic neurological conditions, like stroke, traumatic brain injury, Alzheimer’s disease, Huntington’s disease, and Parkinson’s disease (TBI, AD, HD, and PD, respectively) (Bano et al., 2011; Beal, 1998; Choi, 2020; Hynd et al., 2004).

NADPH-diaphorase neurons [also known as nNOS (+) neurons] are a subpopulation of medium-sized aspiny interneurons, spared following excitotoxic challenges (Granzotto and Sensi, 2015; Koh et al., 1986; Koh and Choi, 1988; Uemura et al., 1990; Weiss et al., 1994). This subpopulation is characterized by the naïve overexpression of the neuronal form of the enzyme nitric oxide synthase (NOS, also known as NOS1) (Dawson et al., 1991; Hope et al., 1991). Notably, early studies have demonstrated nNOS (+) neurons are spared in post-mortem brain samples obtained from AD, HD, and PD patients (Ferrante et al., 1985; Graveland et al., 1985; Mufson and Brandabur, 1994), thereby indicating selective resilience to neurodegeneration.

In two recent studies, we have exploited the distinct features of these neurons to dissect molecular mechanisms associated with the resistance to excitotoxic hits (Canzoniero et al., 2013; Granzotto and Sensi, 2015). These studies have indicated that, in response to NMDA exposures, nNOS (+) neurons produce intracellular Ca^2+^ rises ([Ca^2+^]_i_) that largely overlap with those observed in the general population of nNOS (-) neurons. However, nNOS (+) neurons fail to generate ROS of mitochondrial origin, mobilize neurotoxic amount of Zn^2+^ from intracellular pools and undergo mitochondrial damage (Granzotto et al., 2020).

This study explored distinct subtle differences in NMDAR composition/distribution in nNOS (+) neurons and investigated whether these features offer additional neuroprotective effects. Two lines of evidence support this working hypothesis. First, NMDARs exert different activities according to their subunit composition and synaptic localization (Hardingham and Bading, 2010; Paoletti et al., 2013). Second, it is unclear how nNOS (+) neurons cope with the late-stage (also termed amplification stage) of excitotoxicity, a phase in which damaged neurons spread the toxic cascade to neighboring cells (Zivin and Choi, 1991). Also, not completely clear is the behavior of these cells upon chronic neurodegenerative conditions like AD and HD (Lewerenz and Maher, 2015).

## Materials and methods

### Chemicals

Culture media and sera were purchased from GIBCO (Thermo Fisher Scientific). Fluorescent indicators (fluo-4 AM, fura-2 AM, fura-FF AM) were purchased from Molecular Probes (Thermo Fisher Scientific). NMDA was from Merck Millipore. MK-801 and NBQX were from Alomone. All the other chemicals, unless otherwise stated, were from Sigma-Aldrich.

### Neuronal striatal cultures

All the procedures involving animals were approved by the institutional Ethics Committee (47/2011/CEISA/COM) and carried out following national and international laws and policies. Female mice were caged in groups while male mice were singly housed. Mice were kept on a 12:12 light/dark cycle and had ad libitum access to food and water. All efforts were made to minimize animal suffering during procedures.

Neuronal striatal cultures were prepared as previously described (Granzotto and Sensi, 2015). Briefly, after tissue collection and enzymatic/mechanical dissociation, single-cell striatal suspension was diluted in Neurobasal medium supplemented with 0.5 mM L-glutamine, 5% horse serum, 5% fetal bovine serum, 1 × B27 and 0.2% penicillin/streptomycin and plated onto pre-treated laminin/poly-DL-lysine coated tissue culture plates or dishes. To prevent non-neuronal cell growth and to obtain near-pure striatal cultures, three days after plating, the medium was supplemented with 5 μM of cytosine arabinofuranoside (Ara-C). The 7^th^ day in vitro (DIV) 25% of the medium was replaced with fresh Neurobasal. Experiments were performed on cultures between12 and 18 DIV.

### NADPH-diaphorase staining

NADPH-diaphorase staining was employed for *ex-post* identification of nNOS (+) neurons, as previously described (Canzoniero et al., 2013; Granzotto and Sensi, 2015). Briefly, after microfluorimetry experiments, cells were washed with ice-cold Tris-buffer (TBS), fixed in 4% paraformaldehyde (PFA)/0.1 M phosphate buffer (PBS) for up to 30 minutes at 4° C, rinsed with large volumes of TBS, and incubated for 30-60 minutes at 37° C with freshly made NADPH-diaphorase staining solution containing (in mM): 100 Tris/HCl, 1.2 sodium azide, 0.2 nitrotetrazolium blue, 1 NADPH (Merck-Millipore), and 0.2% Triton X-100, pH 7.2.

### Sample collection for mRNA analysis

Sample collection for mRNA analysis of nNOS (+) and nNOS (-) neurons was performed as described elsewhere (Kim et al., 2001) with some modifications. Striatal cultures were rinsed in ice-cold TBS buffer and fixed in 4% PFA for 15 minutes at 4° C. After PFA removal, cells were thoroughly washed in TBS, permeabilized with Tris/HCl + 0.2 % Triton X100, and stained with the NADPH-diaphorase staining method. The staining solution was then removed, and cells were treated for 20-60 s with Proteinase K (1 μg/ml) in TE buffer (Invitrogen – Thermo Fisher Scientific). Single nNOS (+) neurons were identified on the stage of an upright microscope, aspirated with a patch pipette, and transferred into a 1.5 ml conical tube containing mRNA lysis buffer (PicoPure RNA Isolation Kit -Thermo Fisher Scientific). Similarly, nNOS (-) neurons of similar size and shape were harvested as controls. 11 to 18 neurons per sample were collected from 3 independent cultures. Neurons were lysed and processed following manufacturer instructions and total RNA stored at -80° C until further analysis. All these procedures were performed by employing DEPC water and DNAse/RNAse-free chemicals.

### qRT-PCR analysis

The PicoPure RNA isolation kit (Thermo Fisher Scientific) was employed for total RNA extraction. One µg of RNA was retro-transcribed with the High-Capacity RNA-to-cDNA Kit (Thermo Fisher Scientific). qRT-PCR was carried out on an Abi 7900HT Sequencing Detection System (Thermo Fisher Scientific) in a total volume of 25 μl containing: 2x Maxima SYBR Green/ROX qPCR Master Mix (Thermo Fisher Scientific), 1 µL of cDNA and 0.3 µM of each primer. *Gapdh* and *Hprt1* were employed as endogenous controls. Amplification conditions were as follow: 2 minutes at 50 °C, 10 minutes at 95 °C, followed by 40 cycles of 15 seconds at 95 °C and 1 minute at 60 °C. A melting curve was run to assess the specificity of primers employed. Samples were run in triplicate. The gene relative fold changes were calculated by the ΔΔCt method. The employed primers were: *Gapdh* (Forward 5′-AACAGCAACTCCCACTCTTC-3′, Reverse 5′-GTGGTCCAGGGTTTCTTACTC-3′), *Gpx1* (F 5’-CGACATCGAACCTGACATAGA-3’, R 5’-CAGAGTGCAGCCAGTAATCA-3’), *Grin1* (F 5’-GTGAACGTGTGGAGGAAGAA-3’, R 5’-GTGGAGGTGATAGCCCTAAATG-3’), *Grin2b* (F 5’-GTCCCTTTATCCTCCGTCTTTC-3’, R 5’-CGTCGACTCTCTTGGTTTGTAG-3’), *Grin2a* (F 5’-GCTACTGGAGGGCAACTTATAC-3’, R 5’-TGGTCTGGCAAGAGAGATTTG-3’), *Hprt1* (F 5′-GGCCAGACTTTGTTGGATTTG-3′, R 5′-CGCTCATCTTAGGCTTTGTATTTG-3′), *Nos1* (F 5’-CTCGGTCTTTGTCTCTCTTTCTT-3’, R 5’-GGATGTGATGTGGTAGGGTTAG-3’), *Sod2* (F 5’-GTAGAGCCTTGCCTGTCTTATG-3’, R 5’-AAACCCAGAGGCACCATTAC-3’).

### Bioinformatic analysis

Whole cortex and hippocampus scRNA-Seq data were obtained from the SMART-Seq Allen Brain Atlas database (portal.brain-map.org) and retrieved on March 26^th^, 2020 (Allen Brain Institute, 2019; Lein et al., 2007). nNOS (+) neurons were identified by filtering for neuronal cells showing a high expression of *Nos1* transcripts (>10.0 FPKM) and the abundant expression of additional nNOS(+) neuron markers (*Gad, Sst, Pvalb*, and *Npy*). Two cell clusters (Sst Chodl_1, Sst Chodl_2) were identified (Supplementary File 1). Differentially expressed gene (DEG) analysis was performed using Cytosplore Viewer, a publicly available visual analysis system to interrogate single-cell data published in the Allen Cell Types Database (Tasic et al., 2018). To limit the number of differentially expressed genes and given the heterogeneity of the neuronal subtypes, the nNOS (+)neuron transcriptome was compared with the transcriptome of GABAergic neurons (Supplementary File 2).

### Live-cell imaging

All live-cell imaging experiments were performed on an epifluorescent Zeiss Axio Examiner.D1 upright microscope equipped with a Xenon lamp-based Cairn Optoscan monochromator, a Zeiss 20x NA 1.0 W Plan-Apochromat water immersion objective, and selective fluorescence emission filters. Images were acquired with a Photometrics 16-bit Evolve 512 EMCCD camera and analyzed with the Molecular Devices Metafluor 7.7 software.

### Ca^2+^_i_ imaging experiments

Striatal cultures were loaded for 30 min in the dark at room temperature (RT) with fluo-4 AM (3 µM), fura-2 AM (3 µM), or fura-FF AM (5 µM) plus 0.1% Pluronic F-127 in a HEPES-controlled saline solution (HCSS) containing (in mM): 120 NaCl, 5.4 KCl, 0.8 MgCl_2_, 1.8 CaCl_2_, 20 HEPES, 15 glucose, 10 NaOH, and pH 7.4. Cells were then washed and incubated in the dark for further 30 min in HCSS. fluo-4 (excitation λ: 473 ± 20 nm, emission λ: 525 ± 25 nm) fluorescence changes of each cell were expressed as λF/F, where F is the fluorescence intensity at rest and ΔF the relative fluorescence change (Fx - F) over time. Similarly, fura-2 and fura-FF (excitation λ: 340 ± 10 nm, 380 ± 10 nm, emission λ: 510 ± 45 nm) fluorescence changes of each cell were acquired as 340/380 emission ratio and expressed as ΔR/R, where R is the fluorescence ratio at rest, and ΔR is the relative fluorescence ratio change (Rx - R) over time (Csernansky et al., 1994). During all [Ca^2+^]_i_ measurements, TPEN (200 - 500 nM, Merck Millipore) was added to the bathing solution to prevent interferences of the fluorescent dyes with heavy metal ions (i.e., Zn^2+^) (Grynkiewicz et al., 1985).

### Analysis of spontaneous Ca^2+^_i_ transients

Spontaneous 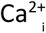 changes were acquired at a 5 Hz sampling rate. Raw fluorescence values of each cell were normalized and analyzed using a cusi tom-made MATLAB script as previously described (Frazzini et al., 2016; Granzotto et al., 2019). The code calculates the number of transients per minute (frequency) and the amplitude of the Ca^2+^ spikes. Only transients that were 50% above the baseline were considered.

### Neuronal striatal culture immunofluorescence

Neuronal striatal cultures were grown on 35 mm glass coverslips. Neurons were washed thoroughly twice in ice-cold PBS, fixed for 10 min at RT with 4% PFA, permeabilized with PBS + 0.1% Triton X-100, and then blocked for 1 h at RT with 1% of bovine serum albumin in PBS + 0.1% Tween-20 (blocking solution). Cells were incubated for 1 h at RT with anti-GluN1 antibody (1:200, Alomone) and anti-NOS1 antibody (1:50, Santa Cruz Biotechnology) in the blocking solution. After washing in PBS cultures were stained with species-specific Alexa-conjugated secondary antibodies (Alexa-633, 1:500; Alexa-488, 1:2000, Thermo Fisher Scientific, respectively) for 1.5 h at room temperature in the dark. Coverslips were then mounted with ProGold-antifade mounting medium (Thermo Fisher Scientific) on cleaned microscopy slides. Cells were imaged on a Zeiss LSM800 confocal microscope equipped with a 63x NA 1.40 Plan-Apochromat oil immersion objective and a super-resolution Airyscan module. Five optical slices (170 nm step size) were acquired for each neuron. After Airyscan processing and background subtraction, images were transformed as maximum orthogonal projections of the whole stack using the ZEN software (Zeiss). Images were further analyzed with the Fiji distribution of ImageJ software as follows. To identify NMDAR-related puncta, GluN1 images were thresholded, the watershed algorithm was applied to define boundaries between the puncta, and finally, binary transformed. The obtained image was used as a mask to measure the number, size, and fluorescent intensity of GluN1 puncta in primarydendrites of nNOS (-) and nNOS (+) neurons.

### Assessment of neuronal injury

Neuronal death was assessed with the lactate dehydrogenase (LDH) efflux assay as previously described (Granzotto and Sensi, 2015).

### Spectroscopic analysis

To evaluate spectroscopic interferences due to DTNB/DTT application, absorbance spectra of the two compounds were measured. DTNB (0.5 mM) and DTT (10 mM) were dissolved in HCSS, and their absorbance spectra measured at room temperature with a SpectraMax 190 plate reader within the 300 – 600 nm range (5 nm step size). Results are reported as optical density (OD).

### Statistical analysis

No statistical methods were employed to determine the sample size. All the results are reported as mean ± standard error of the mean (SEM). Comparison between two groups was performed with Student’s t-test or Welch’s corrected unpaired t-test, where appropriate. For comparisons with more than two groups, one-way or two-way ANOVA was performed, where appropriate, followed by Tukey’s post-hoc test. Based on conventional criteria, results were considered statistically significant when p < 0.05. * indicates p < 0.05 and ** indicates p < 0.01.

## Results

### Spontaneous Ca^2+^ transients of nNOS (+) neurons are identical to the ones of nNOS (-) neurons

Our neuronal cultures exhibit intracellular Ca^2+^ transients that depend on synaptic activity and glutamatergic signaling (Frazzini et al., 2016; Granzotto et al., 2019; Isopi et al., 2015). To evaluate potential differences in the activation of excitatory signaling between nNOS (-) and nNOS (+) neurons, we measured spontaneous Ca^2+^ transients in the two neuronal populations. Striatal neurons were loaded with the high-affinity Ca^2+^ sensitive dye fluo-4 (K_d_ = 335 nM), and changes in [Ca^2+^]_i_ were monitored with microfluorimetry. Changes were analyzed in terms of Ca^2+^ transient frequency and mean transient amplitude. nNOS (-) and nNOS (+) neurons exhibited overlapping patterns of spontaneous activity as far as spiking frequency (Fig. 1A-C) and transient amplitude (Fig. 1D).

**Figure 1.**
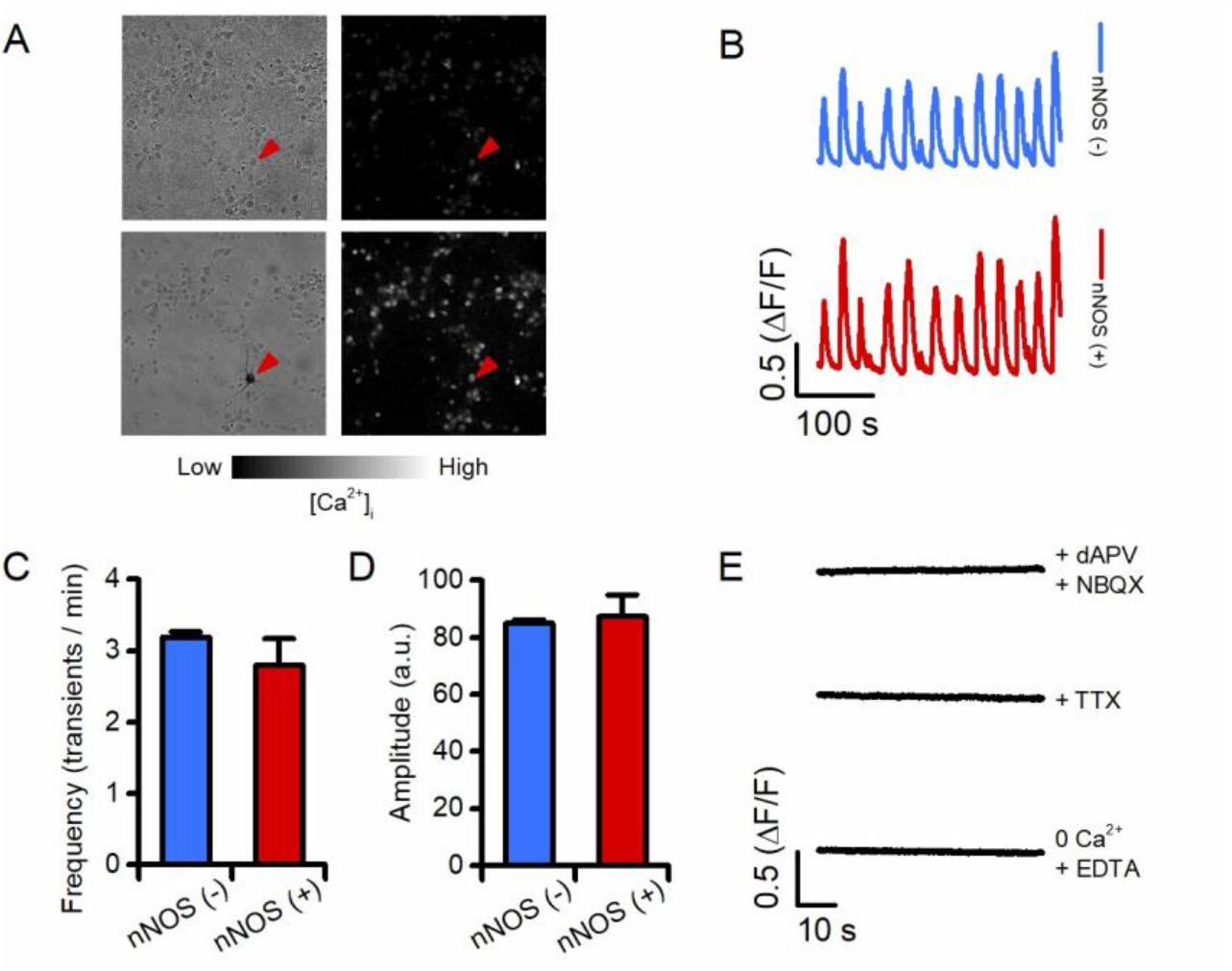
nNOS (+) and the general population of nNOS (-) neurons show overlapping spontaneous Ca^2+^ transients. (A) Striatal neurons were loaded with fluo-4 to monitor spontaneous Ca^2+^ transients *in vitro*. Left panels show phase contrast images of the assayed field before (top) and after (bottom) NADPH-diaphorase staining (cell with dark precipitate, red arrowhead); right panels show greyscale-colored images of fluo-4 loaded cultures before (top) and during (bottom) a Ca^2+^ transient. Images are representative of 7 independent experiments. (B) Representative time courses of Ca^2+^ transients occurring in nNOS (-) and nNOS (+) neurons. (C) Bar graph depicts Ca^2+^ transients frequency values obtained in the two populations (transients/min in nNOS (-): 3.19 ± 0.07 vs. 2.8 ± 0.37 in nNOS (+), p = 0.50, n = 616 nNOS (-) vs. n = 10 nNOS (+) neurons from 7 independent experiments). (D) Bar graph depicts Ca^2+^ transients amplitude values obtained from the same groups as in C (fluo-4 peak amplitude in nNOS (-): 85.01 ± 1.29 vs. 87.39 ± 7.55 in nNOS (+) neurons, p = 0.81). (E) Representative time courses of Ca^2+^ transients occurring in striatal neurons treated with glutamate receptors antagonists (dAPV and NBQX, 100 µM and 10 µM, respectively), TTX (1 µM), or a Ca^2+^-free medium.

Pharmacological manipulations with tetrodotoxin (TTX, 1 μM, to block action potentials), and NBQX (2,3-dihydroxy-6-nitro-7-sulfamoyl-benzo[f]quinoxaline, 10 μM), or dAPV (D-2-amino-5-phosphonovaleric acid, 100 μM) to suppress glutamate-mediated effects demonstrated that changes in [Ca^2+^]_i_ levels are driven by Ca^2+^ entry resulting from action potential firing and activation of synaptic glutamatergic receptors (Fig. 1E). Experiments performed in a Ca^2+^-free medium (supplemented with 50 μM EDTA) fully abrogated [Ca^2+^]_i_ rises (Fig. 1E), thereby indicating that these resulted from Ca^2+^ entry and not mobilization of the cation from intraneuronal sites.

Thus, taken together, these results indicate that nNOS (-) and nNOS (+) neurons do not show significant differences in terms of spontaneous glutamatergic receptor activation.

### Compared to nNOS (-) neurons, nNOS (+) neurons show identical Ca^2+^ rises following synaptic and extrasynaptic NMDAR activation

Several lines of evidence indicate that synaptic and extrasynaptic NMDARs (synNMDARs and exNMDARs, respectively) exert different effects on neuronal functioning, with exNMDARs playing a key role in the activation of pro-death pathways (Hardingham and Bading, 2010).

To evaluate whether the resilience of nNOS (+) neurons to excitotoxic challenges is driven by differences in exNMDARs, we employed an established pharmacological paradigm that selectively and sequentially activates synNMDARs and exNMDARs. After baseline fluorescence acquisition, fluo-4 loaded neurons were exposed to 4-AP (4-aminopyridine, 2.5 mM), a potassium channel blocker. The maneuver promotes sustained neuronal firing, thereby allowing evaluation of Ca^2+^ influx through synaptic glutamatergic receptors (i.e., synNMDARs). Neurons were then exposed to MK-801 (10 μM,in the presence of 4-AP) to promote complete and irreversible blockade of synNMDARs. Following a brief washout period, neurons were then challenged with NMDA (50 μM) + glycine (10 μM) to allow Ca^2+^ entry through exNMDARs.

Analysis of the time course of fluo-4 changes, during synNMDAR activation, revealed no differences in terms of Ca^2+^ rises or cation load between nNOS (-) and nNOS (+) neurons (Fig. 2A-C). Similarly, no significant differences were observed between the two neuronal populations as far as exNMDAR-driven [Ca^2+^]_i_ changes (Fig. 2D-E). Further analysis of [Ca^2+^]_i_ dynamics showed no differences in the cation influx rate (Fig. 2F).

**Figure 2.**
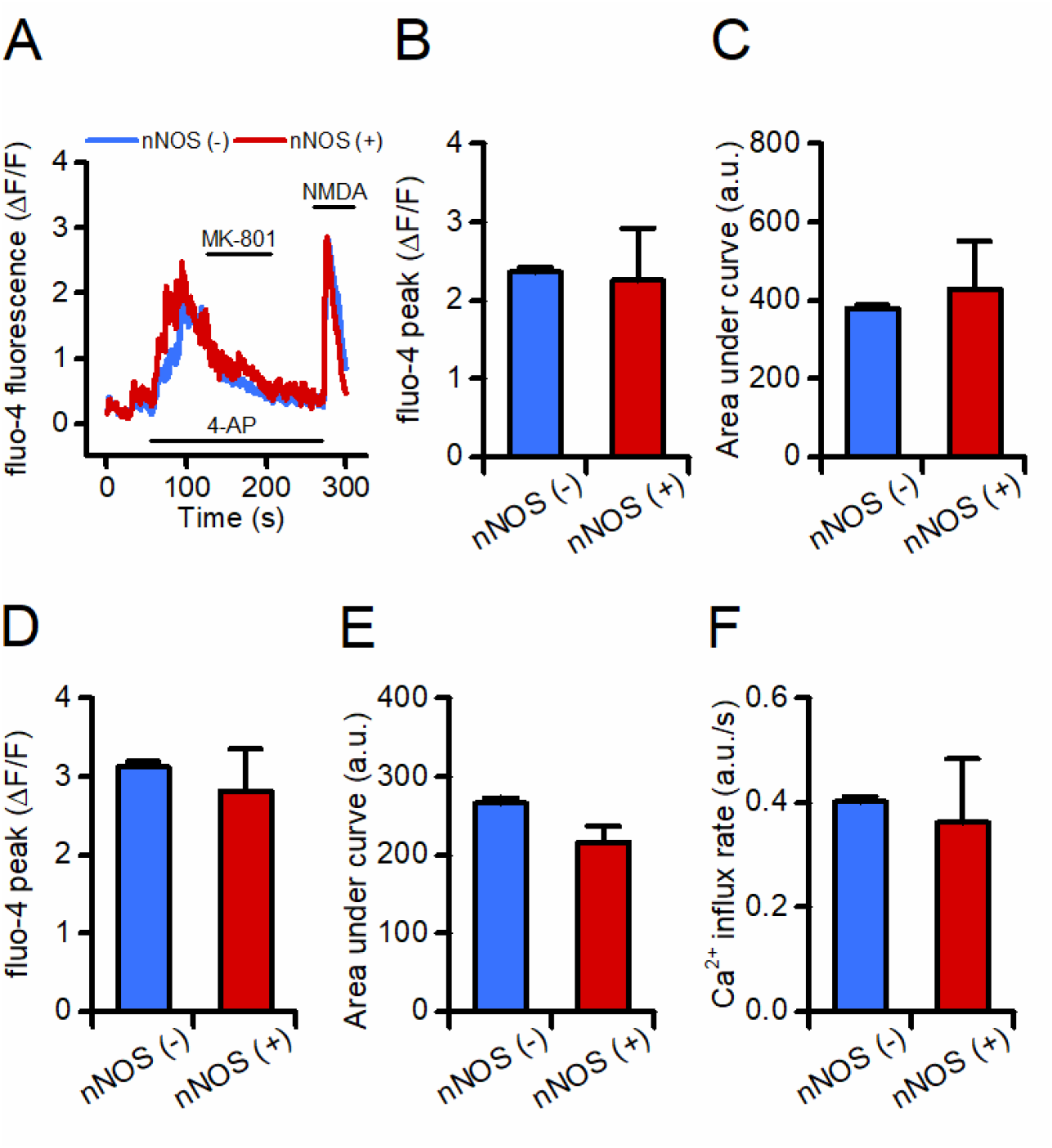
nNOS (+) and the general population of nNOS (-) neurons show overlapping intracellular Ca^2+^ rises upon activation of synaptic and extrasynaptic NMDARs. (A) Representative time courses of fluo-4 loaded nNOS (-) and nNOS (+) striatal neurons exposed to a pharmacological maneuver set to activate synNMDARS and exNMDARs. (B) Bar graph depicts peak of [Ca^2+^]_i_ values obtained in the two populations during synNMDAR activation [fluo-4 peak in nNOS (-): 2.36 ± 0.05 vs. 2.25 ± 0.66 in nNOS (+) neurons, p = 0.81, n = 389 nNOS (-) vs. n = 5 nNOS (+) neurons from 4 independent experiments]. (C) Bar graph depicts cumulative [Ca^2+^]_i_ changes in the two populations expressed as area under the curve (AUC) of arbitrary units (a.u.) during synNMDAR activation [fluo-4 AUC in nNOS (-): 379.28 ± 8.84 vs. 429.01 ± 122.89 in nNOS (+) neurons, p = 0.53]. (D) Bar graph depicts peak of [Ca^2+^]_i_ values obtained in the two populations during exNMDAR activation [fluo-4 peak in nNOS (-): 3.12 ± 0.06 vs. 2.81 ± 0.53 in nNOS (+) neurons, p = 0.59]. (E) Bar graph depicts cumulative [Ca^2+^]_i_ changes in the two populations during exNMDAR activation [fluo-4 AUC in nNOS (-): 266.08 ± 7.33vs. 216.22 ± 20.61 in nNOS (+) neurons, p = 0.07]. (F) Bar graph depicts Ca^2+^ influx rate in the two populations expressed as a.u. changes per second during exNMDAR activation (influx rate in nNOS (-) 0.40 ± 0.009 vs. 0.36 ± 0.12 in nNOS (+) neurons, p = 0.63].

Overall, these findings indicate that nNOS (+) and (-) neurons respond to the activation of synNMDARs and exNMDARs with overlapping changes in [Ca^2+^]_i_, thereby suggesting that the two populations are equipped with similar pools of equally functional synNMDARs and exNMDARs.

### Compared to nNOS (-) neurons, nNOS (+) neurons show reduced transcriptomic and protein expression of the NMDAR subunit 1

Previous studies investigating the expression of NMDAR subunits in nNOS (+) neurons have provided contrasting results (Augood et al., 1994; Kim et al., 2001; Landwehrmeyer et al., 1995; Price Jr. et al.,1993; Weiss et al., 1998). To address this question in our system, we performed qRT-PCR on a set of selected transcripts obtained from pools of nNOS (-) and nNOS (+) neurons (11 to 18 neurons per sample, Fig. 3A). Analysis of qRT-PCR data showed that, compared to nNOS (-), nNOS (+) neurons exhibit reduced expression of *Grin1*, the gene encoding for the mandatory NMDAR subunit GluN1 (also known as NR1) (Fig. 3B). Other NMDAR candidate gene transcripts, *Grin2a* and *Grin2b*, show no differences between the two populations (Fig. 3B). Expression levels of other genes coding for proteins that have been proposed to be involved in the neuroprotection exhibited by nNOS (+) neurons, like *Sod2* and *Gpx1*, were similar in the two study groups. The *Bcl2* transcript was not detectable (Fig. 3B and Supplementary Table 1). Of note, *Nos1* transcript (encoding the nNOS protein) was found significantly increased in nNOS (+) neuron samples, thereby confirming the selectivity and specificity of our procedure in the isolation of nNOS (+) neurons (Fig. 3B).

**Figure 3.**
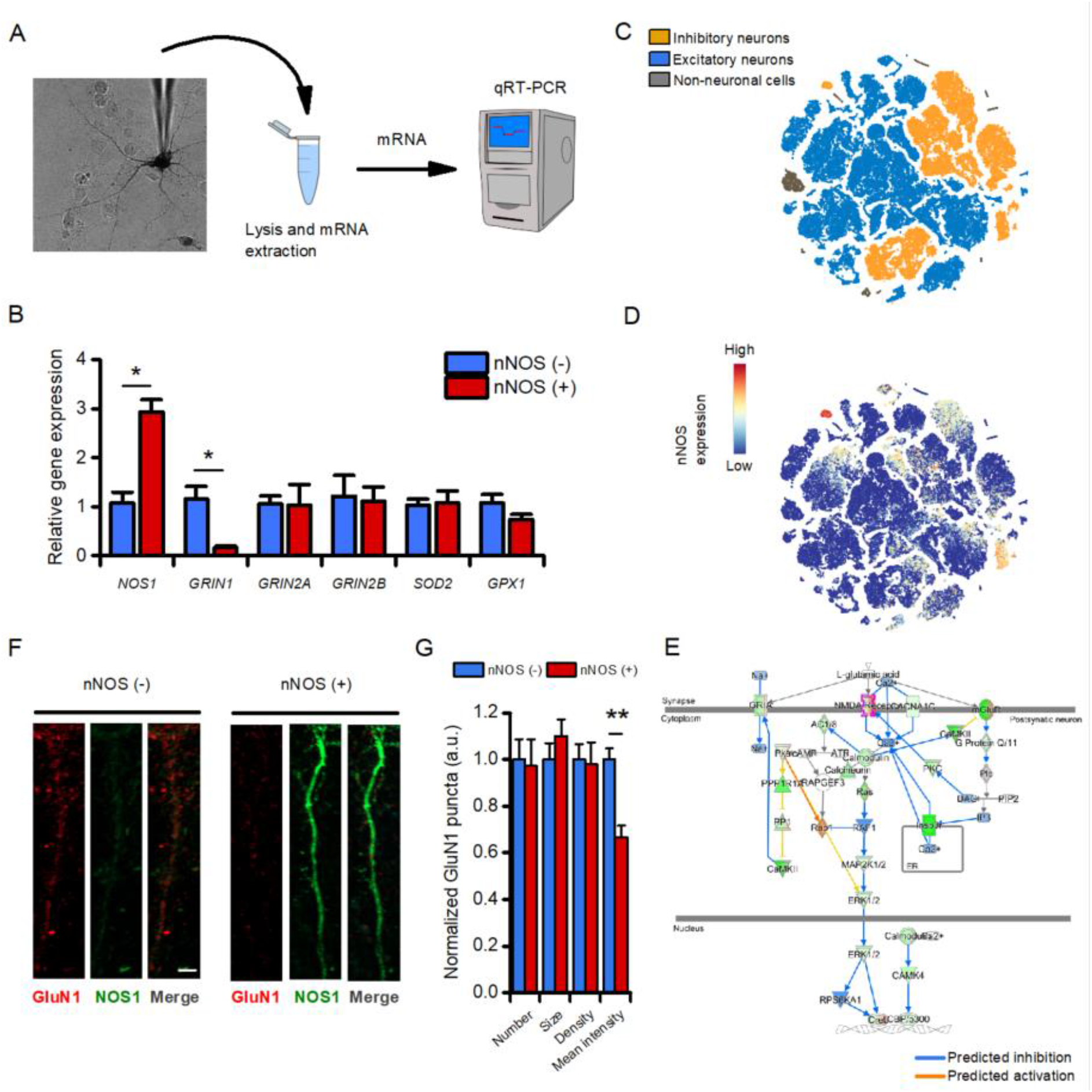
nNOS (+) neurons show a decreased expression of the NMDA receptor subunit GluN1. (A) The pictogram illustrates the “patch-like” procedure employed to isolate mRNA obtained from single nNOS (+) neurons. The same approach was employed to isolate nNOS (-) neurons employed as control. (B) Bar graphs illustrate changes in mRNA levels of the indicated genes measured by real-time PCR in nNOS (+) neurons when compared to the general population of nNOS (-) cells (relative *Nos1* expression in nNOS (-): 1.07 ± 0.22 vs. 2.92± 0.25 in nNOS (+) neurons, p < 0.001; relative *Grin1* expression in nNOS (-): 1.15 ± 0.25 vs. 0.16 ± 0.02 in nNOS (+) neurons, p = 0.005; n = 4-5 replicates). No differences were observed in the other tested transcripts (see Supplementary Table 1). (C) t-SNE plot of the scRNA-Seq database from the Allen Brain Atlas. Clusters are color-coded basedon cell classification as inhibitory neurons (yellow), excitatory neurons (blue), or non-neuronal cells (gray). (D) t-SNE plot of the same dataset as in C showing levels of nNOS expression. Please, note the presence of a high nNOS expressing cluster (top left) identified as *bona fide* nNOS (+) neurons. (E) The pictogram illustrates the glutamatergic signaling Pathway Activity Analysis. The analysis, based on the expression of significantly perturbed genes from our dataset, predicts if the pathway is activated (yellow-orange arrows) or inhibited (blue arrows) in nNOS (+) neurons when compared to excitatory neurons. Please, note the consistent inhibition of glutamatergic signaling. (F) Representative super-resolution confocal images of dendrites obtained from striatal nNOS (-) (left panel) and nNOS (+) neurons and stained with anti-GluN1 (red) and anti-NOS1 (green) antibodies (scale bar = 2 µm). (G) Bar graph depicts quantification of dendritic GluN1-related fluorescent intensity [normalized GluN1 signal in nNOS (-): 1.00 ± 0.04 vs. 0.66 ± 0.05 in nNOS (+) neurons, p<0.001, n = 43 – 44 dendrites from at least 3 independent experiments].

Our transcriptomic findings were further validated by interrogating mouse scRNA-Seq data obtained from the Allen Brain Cell Atlas database. We identified two neuronal clusters expressing specific nNOS (+) markers (*Nos1*+, *Sst*+, *Gad*+, *Pvalb*+, and *Npy*+; Fig. 3C-D and Supplementary File 1). DEGs assessment was performed by comparing the transcriptome of nNOS (+) neurons with the transcriptome of the broad family of excitatory neurons (Supplementary File 2), a population particularly vulnerable in neurodegenerative settings (Fu et al., 2018). In line with the aim of the study, the downstream analysis focused on elucidating differences in terms of glutamatergic signaling. Supporting our qRT-PCR results, gene ontology (GO) analysis showed that nNOS (+) neurons display an overall reduction of transcripts associated with excitatory glutamatergic signaling (Fig. 3E and Supplementary File 2). In line with transcriptomic data, GluN1 reduction was confirmed by immunofluorescence (IF) in cultured striatal neurons (Fig. 3F-G).

### Compared to nNOS (-), nNOS (+) neurons show reduced NMDAR-driven [Ca^2+^]_i_ rises following receptor reduction

NMDAR ionic conductance can be modulated by oxidizing/reducing agents (Aizenman et al., 1989, 1990, 2020). We evaluated, in nNOS (+) and nNOS (-) neurons, NMDAR activity before and after pharmacological manipulation set to alter the redox status of the receptor (Fig. 4A). After baseline fluorescence acquisition, fura-2 loaded neurons were challenged with NMDA (25 μM) + glycine (2-5μM). After agonist washout, cells were sequentially exposed to 5,5′-dithiobis(2-nitrobenzoic acid) (DTNB; 0.5 mM) and to dithiothreitol (DTT; 2-10 mM), and finally to NMDA (25 μM) + glycine (2-5μM). The maneuver allows the evaluation of NMDAR activity in conditions of receptor reduction state (Aizenman et al., 1989). Results were analyzed in fold changes of NMDA-driven Ca^2+^_i_ rises before and after DTNB-DTT exposures. While DTNB-DTT application resulted in a net increase of NMDA-driven Ca^2+^_i_ entry in nNOS (-) neurons (Fig. 4B-C), the same maneuver produced significantly lower Ca^2+^_i_ changes in nNOS (+) neurons (Fig. 4B-C). Similar results were observed when cultures were exposed to a higher concentration of DTT (10 mM; Fig. 4C). Of note, DTNB-DTT application did not modify resting Ca^2+^_i_ levels in the two populations (Fig. 4D), thereby indicating that fura-2 signals are not affected by artifactual differences in resting levels of cation load.

**Figure 4.**
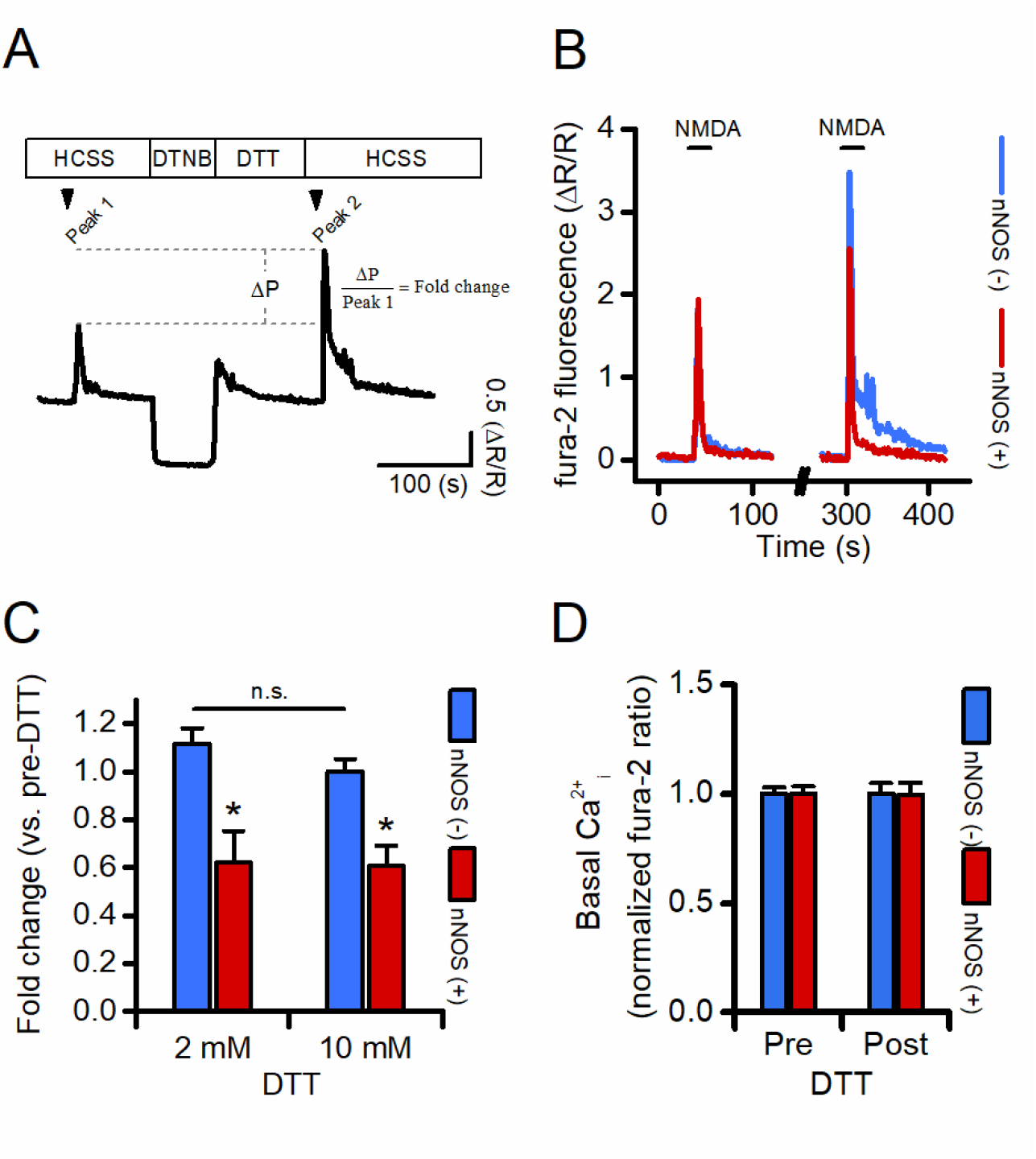
nNOS (+) neurons show decreased NMDAR-dependent Ca^2+^ rises following pharmacological receptor reduction. (A) The pictogram illustrates the pharmacological protocol set to evaluate agonist-dependent changes in [Ca^2+^]_i_ rises before and after NMDAR oxidation/reduction. Please, note that the sharp decrease in fura-2 signal upon DTNB exposure is due to spectroscopic interferences between the probe and the drug (Supplementary fig. 1). (B) Representative time courses of fura-2 loaded nNOS (-) and nNOS (+) striatal neurons exposed to NMDA before (left traces) and after (right traces) full receptor reduction (for clarity, traces during DTNB and DTT exposure were omitted). (C) Bar graphs depict quantification of experiments in B expressed as fold changes in [Ca^2+^]_i_ rises following exposure to 2 mM (left panel) or 10 mM (right panel) DTT in the two populations (DTT 2mM, fura-2 fold change in nNOS: (-) 1.11 ± 0.06 vs. 0.61 ± 0.13 in nNOS (+) neurons, p = 0.005, n = 389 nNOS (-) vs. n = 9 nNOS (+) neurons from 7 independent experiments; 10 mM, fura-2 fold change in nNOS (-): 0.99 ± 0.05 vs. 0.60 ± 0.08 in nNOS (+) neurons, p = 0.002, n = 342 nNOS (-) vs. n = 7 nNOS (+) neurons from 5 independent experiments). (D) Bar graph depicts basal Ca^2+^_i_ levels in nNOS (-) and nNOS (+) neurons before and after DTT exposure expressed as normalized fura-2 ratio. n.s. = not significant.

Overall, this set of experiments supports the notion that nNOS (+) neurons possess a larger pool of fully reduced NMDARs when compared to the general population of nNOS (-) neurons.

### Long-term dynamic changes in NMDAR levels could account for nNOS (+) neurons survival following an excitotoxic challenge

nNOS (+) neurons are spared from acute excitotoxic insults by occluding critical steps in the cascade (Canzoniero et al., 2013; Granzotto and Sensi, 2015; Granzotto et al., 2020). To evaluate additional mechanisms of neuroprotection, we set a protocol that mimics long-term excitotoxic insults and imaged functional changes in nNOS (+) neurons.

Serum- and supplement-free Neurobasal medium exchange has been found to produce, in mature, near-pure cultured neurons, widespread neuronal loss, a phenomenon primarily driven by L-cysteine-dependent activation of NMDARs (Hogins et al., 2011; Maggioni et al., 2015; Olney et al., 1990).

In line with a previous report (Hogins et al., 2011), we found that medium exchange resulted in 40% neuronal loss in our cultures 16 to 24 hours after the challenge (Fig. 5A-B). The toxic effect was abolished when the medium exchange was performed in the presence of dAPV (100 µM). Of note, nNOS (+) neurons were largely spared from the maneuver (Fig. 5A, C and unpublished observations).

**Figure 5.**
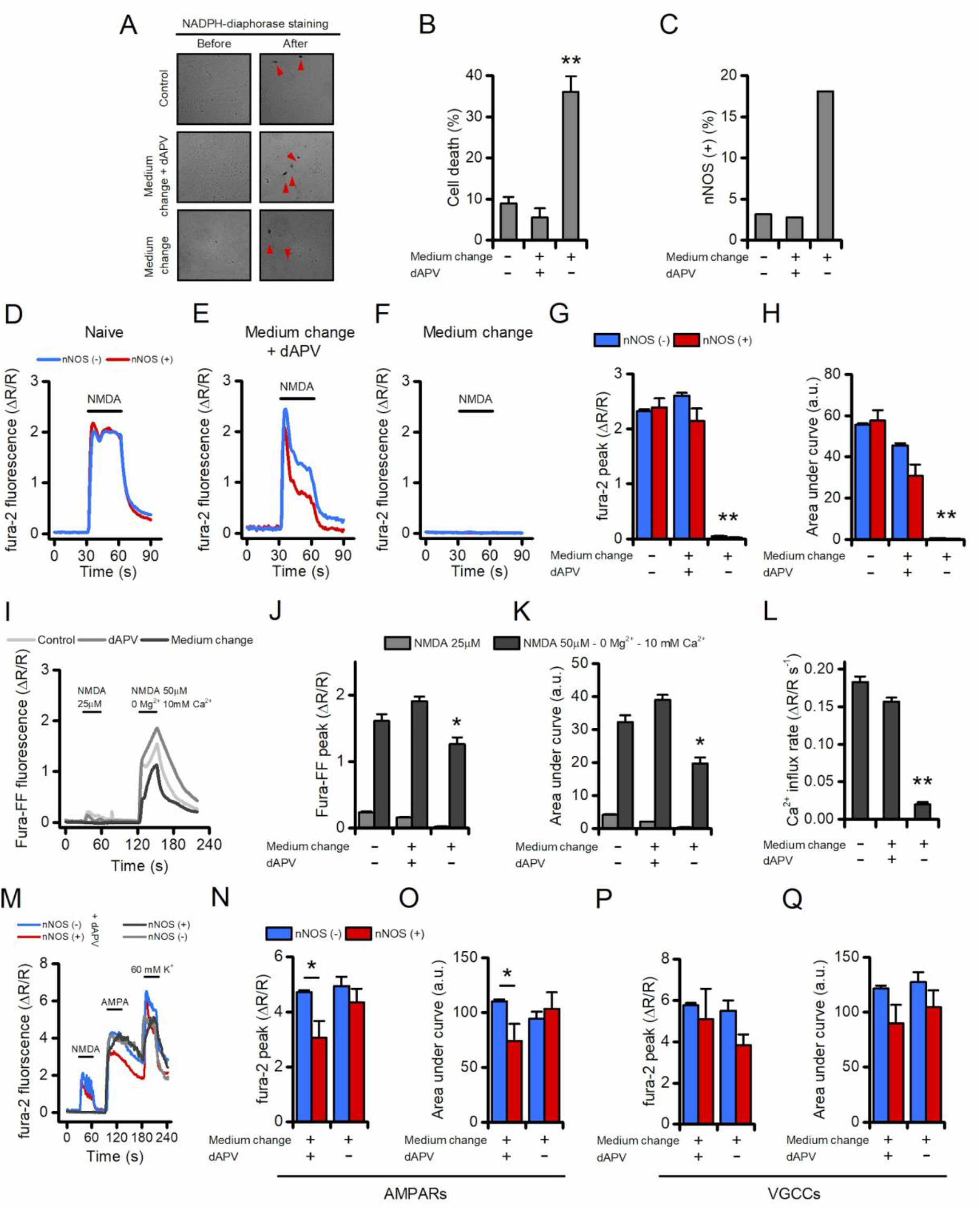
Neurons spared after an excitotoxic challenge fail to respond to NMDA stimulation. (A) The pictogram illustrates phase contrast images of untreated (top), medium change + dAPV-treated (middle), and medium change-treated (bottom) neuronal striatal cultures before (left) and after (right) the NADPH-diaphorase staining. Red arrowheads indicate nNOS (+) neurons. (B) Bar graph depicts the vulnerability of striatal cultures exposed to the treatments described in A. Neuronal viability was assessed, with LDH efflux assay, 16 h after the challenge (neuronal death in naïve neurons: 8.9 ± 1.5 % vs. 5.4 ± 2.2 % in dAPV group vs. 36.0 ± 3.8 % in medium exchange group, F(2, 45) = 37.56, p < 0.0001). (C) Bar graph depicts the increased number of nNOS (+) neurons, expressed as % of the live neurons, following medium exchange treatment which is indicative of a relative sparing of the subpopulation. (D-F) Representative time courses of fura-2 loaded nNOS (-) and nNOS (+) striatal neurons exposed to NMDA (25 µM + 2-5 µM glycine) 16-20 h after being exposed to the indicated treatment. (G) Bar graphs show quantification of fura-2 peak values obtained from experiments shown in D-F (treatment effect F_(2, 1510)_ = 119.7, p < 0.0001; cell type effect F_(2, 1510)_ = 0.9163, p = 0.33; interaction F_(2, 1510)_ = 1.357, p = 0.25). (H) Bar graphs show quantification of cumulative [Ca^2+^]_i_ changes obtained from experiments shown in D-F (treatment effect F_(2, 1510)_ = 107.7, p < 0.0001; cell type effect F_(2, 1510)_ = 1.793, p = 0.18; interaction F_(2, 1510)_ = 2.690, p = 0.07). (I) Representative time courses of fura-FF loaded neuronal striatal cultures exposed to 25 µM NMDA (2-5 µM glycine) or 50 µM NMDA (10 µM glycine) in a Mg^2+^-free medium supplemented with 10 mM CaCl_2_ and assessed 16-20 h after being challenged with the indicated treatment. (J) Bar graph shows quantification of fura-FF peak values obtained from experiments shown in I (F_(2, 402)_ = 16.11, p < 0.0001). (K) Bar graphs show quantification of cumulative [Ca^2+^]_i_ changes obtained from experimentsshown in I (F_(2, 402)_ = 33.43, p < 0.0001). (L) Bar graph depicts Ca^2+^ influx rate in the three treatment groups expressed as a.u. changes per second during the first 5 s of the 50 µM NMDA stimulation (F_(2, 402)_ = 209.2, p < 0.0001). (M) Representative time courses of fura-2 loaded nNOS (-) and nNOS (+) striatal neurons sequentially exposed to NMDA (25 µM + 2-5 µM glycine), AMPA (100 µM + cyclothiazide), or a high K^+^ solution (60 mM K^+^, 10 µM MK-801, 10 µM NBQX). (N) Bar graph shows quantificationof fura-2 peak values obtained from neurons exposed to AMPA (treatment effect F_(1, 459)_ = 2.889, p = 0.09; cell type effect F_(1, 459)_ = 6.383, p = 0.01; interaction F_(1, 459)_ = 1.382, p = 0.24). (O) Bar graph shows quantification of cumulative [Ca^2+^]_i_ changes obtained from neurons exposed to AMPA (treatment effect F_(1, 459)_ = 0.4273, p = 0.51; cell type effect F_(1, 459)_ = 1.622, p = 0.20; interaction F_(1, 459)_ = 4.577, p = 0.03). (P) Bar graph shows quantification of fura-2 peak values obtained from neurons exposed to ahigh K^+^ solution (treatment effect F_(1, 459)_ = 1.137, p = 0.28; cell type effect F_(1, 459)_ = 2.676, p = 0.10; interaction F_(1, 459)_ = 0.4707, p = 0.49). (Q) Bar graph shows quantification of cumulative [Ca^2+^]_i_ changes obtained from neurons exposed to a high K^+^ solution (treatment effect F_(1, 459)_ = 0.4063, p =0.52; cell type effect F_(1, 459)_ = 2.760, p = 0.09; interaction F_(1, 459)_ = 0.073, p = 0.78).

Thus, we employed the neurobasal medium change maneuver to dissect changes in cells exposed to a long-term excitotoxic environment (Hogins et al., 2011; Olney et al.,1990). Neurobasal-treated cultures were compared with sister cultures challenged in the presence of dAPV. Cultures with no medium exchange (naïve cells) were used as control. Sixteen to twenty hours after the insult cultures were loaded with fura-2 and NMDA-driven Ca^2+^_i_ changes evaluated. nNOS (+) and few surviving nNOS (-) neurons did not show alteration in [Ca^2+^]_i_ while dAPV-treated and naïve sister cultures produced significant [Ca^2+^]_i_ elevations (Fig. 5C-G). Of note, naïve and dAPV-treated cultures showed differences in overall cation load (Fig. 5H), thereby suggesting that, in our model, dAPV exposures positively affect Ca^2+^ buffering mechanisms.

To test if the abolished response to NMDA depends on reduced expression or defective functioning of the receptor, we exposed our cultures to a pharmacological maneuver that triggers maximal NMDARs activity. To assess these large Ca^2+^_i_ changes, the low-affinity Ca^2+^ sensor fura-FF (K_d_ = 5.5 µM) was used, and cultures were challenged with NMDA (50 μM) + glycine (10 μM) in a magnesium-free medium supplemented with 10 mM Ca^2+^. In line with experiments shown in Fig. 5C-E, medium exchange-treated cultures exhibited significantly reduced [Ca^2+^]_i_ amplitudes, overall cation loads, and cation influx rates compared to the control groups (Fig. 5I-L).

Exposures to AMPA or a depolarizing medium (high K+, Fig. 5M-Q) failed to generate Ca^2+^_I_ rises, thereby indicating that Ca^2+^_i_ changes were specifically driven by NMDAR activation (Turetsky et al., 1994).

## Discussion

The primary purposes of the study were to 1) elucidate the molecular and functional properties of NMDARs in the subpopulation of nNOS (+) neurons and 2) evaluate effects of chronic excitotoxic challenges. Our results integrate our previous findings (Canzoniero et al., 2013; Granzotto and Sensi, 2015) and provide novel insights on the role of NMDARs in nNOS (+) neurons and may help decipher the role of NMDARs under neurodegenerative conditions. Our study supports the notion of dynamic control of NMDAR activity at least in part modulated by the neuronal redox status and the presence of a toxic extracellular milieu.

NMDAR-driven intracellular Ca^2+^ overload is a mandatory step in the excitotoxic cascade. Compelling evidence, however, suggests that receptor subunit arrangements and localization significantly affect the downstream responses elicited by NMDAR agonists, independently of overall cation accumulation.

The activity of the NMDAR is affected by its subunits (Paoletti et al., 2013). Endogenous modulators like protons, Zn^2+^, and ROS exert an inhibitory effect (Zhu and Paoletti, 2015) while specific amino acids (glycine and D-serine) or reducing agents potentiate NMDAR activity (Paoletti et al., 2013). The complexity of NMDARs physiology is exacerbated by the divergent action played by synNMDARs and exNMDARs. Evidence indicates that exNMDARs promote cell death signaling in antagonism to synNMDARs that activate anti-apoptotic and neurotrophic pathways (Hardingham et al., 2002; Hardingham and Bading, 2010). We combined functional, transcriptomic,and imaging data to gain further insights into the role of NMDAR activation in the nNOS (+) neurons resilience to excitotoxins.

Our results revealed a composite picture. Ca^2+^ imaging experiments, in line with our previous reports (Canzoniero et al., 2013; Granzotto and Sensi, 2015), indicate that nNOS (+) neurons possess equally functional pools of NMDARs when compared to nNOS (-) neurons. In addition, no topological differences were observed when evaluating synNMDARs and exNMDARs activity in the two neuronal populations (Fig. 2). Surprisingly, the transcriptomic and histochemical analysis indicated a net decrease in the expression of NMDA GluN1 subunit (Fig. 3) in nNOS (+) neurons, an important feature considering that, among the seven different NMDAR subunits, GluN1 plays a mandatory role for the receptor assembling and functioning (Paoletti et al., 2013).

To gain some further insight, we focused on two distinguishing features of the NMDARs and the nNOS (+) subpopulation. NMDARs are modulated by oxidizing and reducing agents, which decrease or potentiate the amplitude of receptor response to agonists, respectively (Aizenman et al.,1989, 1990, 2020). On the other hand, nNOS (+) neurons fail to generate ROS of mitochondrial origin following the activation of NMDARs (Canzoniero et al., 2013; Granzotto and Sensi, 2015). The phenomenon may be related to increased cellular defenses against oxidative/nitrosative damage (Gonzalez-Zulueta et al., 1998; Granzotto and Sensi, 2015; Granzotto et al., 2020). This distinct feature supports the idea that nNOS (+) neurons, by constitutively dealing with an antioxidant milieu, possess fewer but more reduced, and therefore more functional, NMDARs. This notion is supported by findings on NMDAR activity in striatal neurons exposed to DNTB/DTT (Fig. 4). This set of experiments indicates that, upon complete NMDAR reduction, nNOS (+) neurons generate significantly reduced receptor-mediated Ca^2+^_i_ amplitudes when compared to nNOS (-) neurons (Figs. 4C). Thus, one can speculate that, in nNOS (+) neurons, a cell-autonomous mechanism regulates *GluN1* expression to balance increased receptor functioning. This intriguing hypothesis is supported by complementary evidence showing that non-toxic oxidative challenges, which conceivably reduce NMDAR activity (Aizenman et al., 1990), result in upregulated *GluN1* expression (Betzen et al., 2009; Hota et al., 2010; Massaad and Klann, 2011). In addition, this view extends recent findings on the mechanistic liaison between NMDAR activity and the transcription of antioxidant molecules (Baxter et al., 2015; Papadia et al.,2008) and open the possibility for a feedback loop in which the cell redox status may affect the transcription of synaptic proteins and vice versa. This hypothesis may also have implications for disorders in which NMDAR overactivation, increased ROS generation, and alterations in the transcriptional machinery help to modulate neurodegenerative processes.

Moreover, with the limitations of an indirect, population-based approach, our results confirm that nNOS (+) neurons possess fully operational NMDARs [Figs. 1 and 2 and (Canzoniero et al., 2013; Granzotto and Sensi, 2015)], thereby arguing against the presence of mechanisms of resistance that act upstream in the excitotoxic cascade. We also found that changes in NMDARs may account for nNOS (+) resilience upon chronic excitotoxic hits. This idea is supported by a set of experiments showing that nNOS (+) neurons fail to respond to NMDA stimulations after prior exposure to chronic excitotoxic challenges (Fig. 5). The effect was found to be specific to NMDAR- and not AMPAR-or VGCC-dependent activation and mirrors the attenuation of NMDAR activity previously reported in an *in vivo* model of TBI (Biegon et al., 2004). In this regard, one can speculate that nNOS (+) neurons, by missing critical steps of the early stages of the excitotoxic cascade (Canzoniero et al., 2013; Granzotto and Sensi, 2015) activate pathways instrumental for NMDA removal in the attempt to limit the damage associated with chronic excitotoxicity.

Three major evidence supports this idea. First, NMDARs, although reported to be static when compared to AMPARs, possess endocytic motifs that are required for receptor internalization and degradation (Roche et al., 2001; Scott et al., 2004). Second, when stimulated to operate at full capacity, NMDARs generated lower [Ca^2+^]_i_ rises (Figs. 5I-L) when compared to control cultures, thereby suggesting that fewer receptors are present on the cellular surface. The third argument is specific to our experimental setting as the activation of the NMDAR glycine (and L-cysteine) binding site primes the receptor internalization (Nong et al., 2003). In agreement, excitotoxic challenges performed in the absence of NMDAR co-agonists (i.e., glycine or D-serine) produce different functional and viability outcomes (Wu et al., 2017). Of note, this proposed mechanism is not limited to nNOS (+) neurons but can be extended to virtually all those neurons that are spared by our chronic excitotoxic challenge (Fig. 5A-B). However, we cannot exclude the possibility that these subsets of neurons fail to respond to NMDA stimulations for reasons unrelated to receptor expression on the plasma membrane (i.e., negative post-translational modifications, etc.). Another unsolved question is whether the blockade of NMDAR signaling is a regulated process. Further studies aimed at manipulating the underlying mechanism will be required.

## Conclusions

Our findings may have intriguing implications for neurological conditions associated with NMDAR overactivation. The reduced vulnerability of nNOS (+) neurons indicates that the presence of downstream steps of the cascade can be promising pharmacological targets for neuroprotection (Granzotto et al., 2020). The phenomenon also can limit the side effects associated with the pharmacological blockade of NMDARs (Ikonomidou and Turski, 2002). In addition, the reduced NMDAR responses following chronic excitotoxic hits provide alternative heuristic models to understand the failure of NMDAR antagonists in clinical trials as well as gain insight into the mechanisms associated with the neuroprotective effects exerted by preconditioning against ischemic neuronal death (Aizenman et al., 2000; Dirnagl et al., 2009).

## Author contributions

Conceptualization, Alberto Granzotto and Stefano Sensi; Formal analysis, Alberto Granzotto and Valentina Gatta; Funding acquisition, Alberto Granzotto, Valentina Gatta and Stefano Sensi; Investigation, Alberto Granzotto, Marco d’Aurora and Manuela Bomba; Methodology, Alberto Granzotto and Valentina Gatta; Project administration, Stefano Sensi; Visualization, Alberto Granzotto; Writing – original draft, Alberto Granzotto; Writing – review & editing, Alberto Granzotto, Valentina Gatta, Marco Onofrj and Stefano Sensi.

## Acknowledgments

The authors thank all the members of the Molecular Neurology Unit for helpful discussions.

---

**Supplementary table 1.**

| **Gene** | **nNOS (-)** | **nNOS (+)** | ***P*** |
| --- | --- | --- | --- |
| ***Nos1*** | 1.07 ± 0.22 | 2.92 ± 0.25 | 0.0003 |
| ***Grin1*** | 1.15 ± 0.25 | 0.16 ± 0.02 | 0.0272 |
| ***Grin2a*** | 1.05 ± 0.16 | 1.02 ± 0.42 | 0.31 |
| ***Grin2b*** | 1.21 ± 0.42 | 1.11 ± 0.29 | 0.42 |
| ***Sod2*** | 1.03 ± 0.12 | 1.07 ± 0.24 | 0.42 |
| ***Gpx1*** | 1.06 ± 0.17 | 0.73 ± 0.1 | 0.08 |
| ***Bcl2*** | ND | ND |  |

**Supplementary figure 1.**
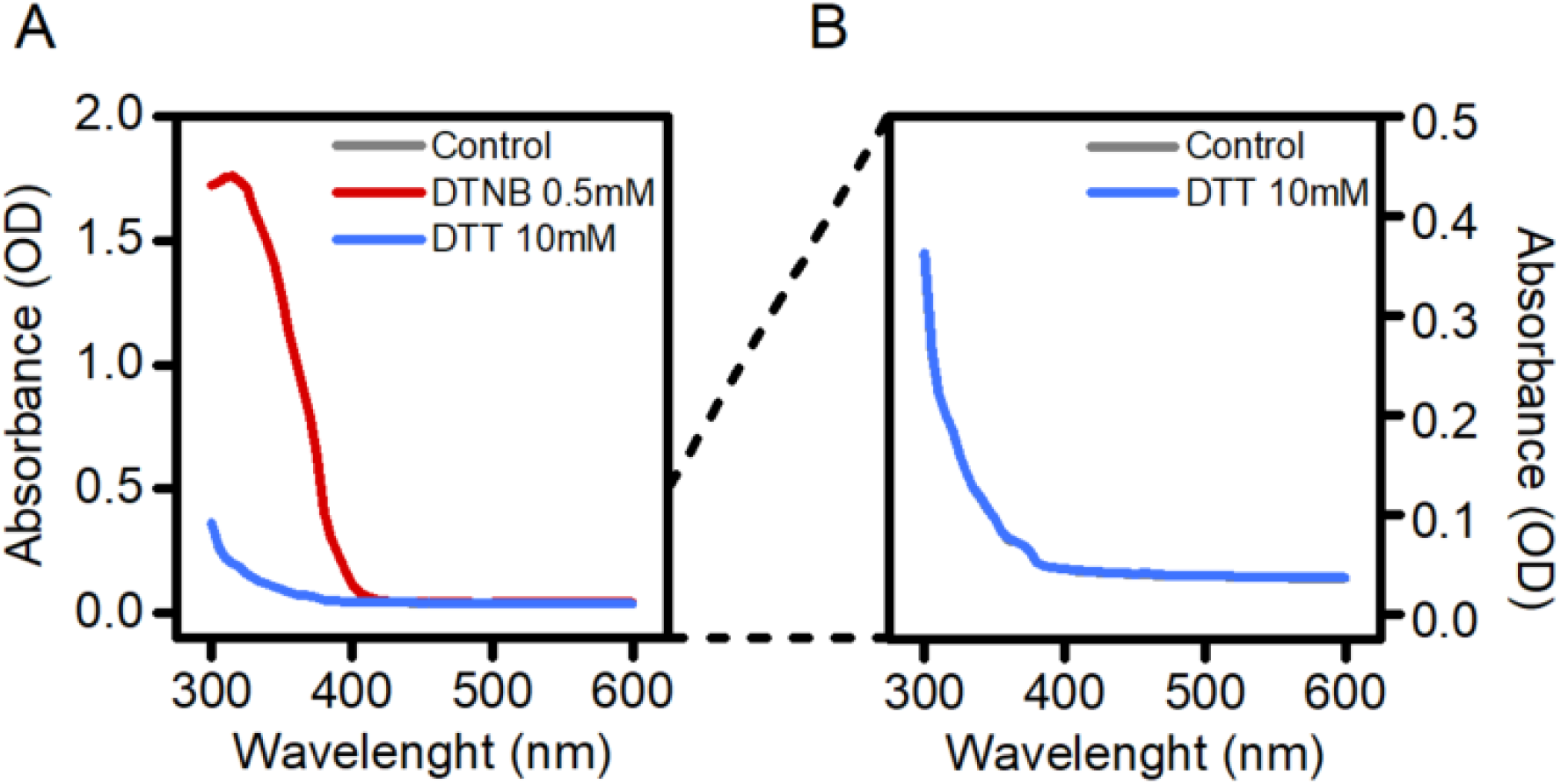
DTNB interferes with fura-2 excitation wavelengths. (A) The pictogram illustrates DTNB (0.5 mM) and DTT (10 mM) absorption spectra recorded in the 300 – 600 nm range (5 nm step size). Please note that DTNB shows maximum absorbance at 340 nm thereby interfering with the short fura-2 excitation wavelength (340/380 nm). (B) The pictogram shows a magnification of DTT spectrum. Please, note the complete overlap between DTT and control medium traces.

