## Supplementary figures and images for "Long-term dynamic changes of NMDA receptors following an excitotoxic challenge"

### Supplementary Figure 1

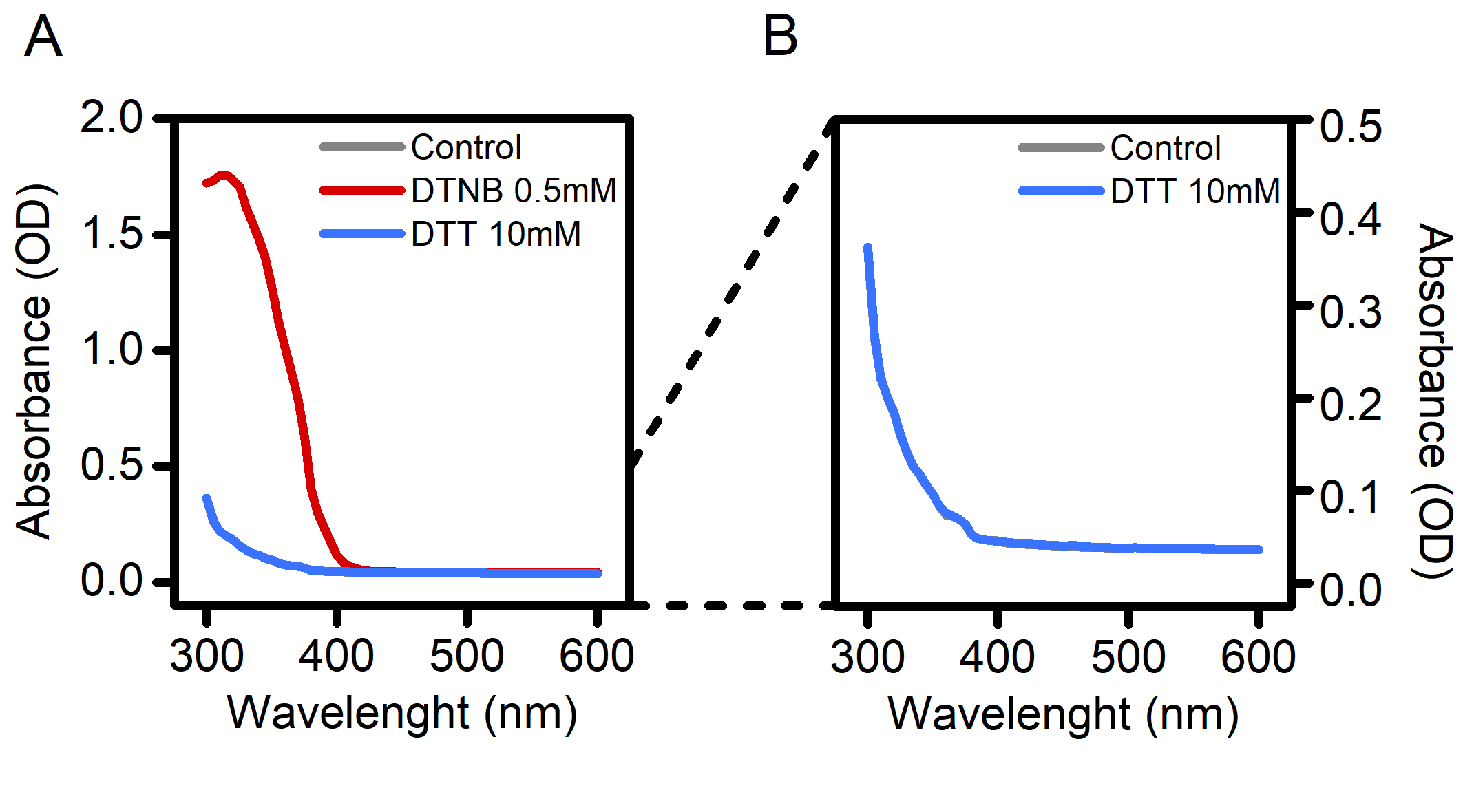
